# Heterochromatin delays CRISPR-Cas9 mutagenesis but does not influence repair outcome

**DOI:** 10.1101/267690

**Authors:** Eirini M Kallimasioti-Pazi, Keerthi Thelakkad Chathoth, Gillian C Taylor, Alison Meynert, Tracy Ballinger, Martijn Kelder, Sébastien Lalevée, Ildem Sanli, Robert Feil, Andrew J Wood

## Abstract

CRISPR-Cas9 genome editing occurs in the context of chromatin, which is heterogeneous in structure and function across the genome. Chromatin heterogeneity is thought to affect genome editing efficiency, but this has been challenging to quantify due to the presence of confounding variables. Here, we develop a method that exploits the allele-specific chromatin status of imprinted genes in order to address this problem. Because maternal and paternal alleles of imprinted genes have identical DNA sequence and are situated in the same nucleus, allele-specific differences in the frequency and spectrum of Cas9-induced mutations can be attributed unequivocally to epigenetic mechanisms. We found that heterochromatin can impede mutagenesis, but to a degree that depends on other key experimental parameters. Mutagenesis was impeded by up to 7-fold when Cas9 exposure was brief and when intracellular Cas9 expression was low. Surprisingly, the outcome of mutagenic DNA repair was independent of chromatin state, with similar efficiencies of homology directed repair and deletion spectra on maternal and paternal chromosomes. Combined, our data show that heterochromatin imposes a permeable barrier that influences the kinetics, but not the endpoint of CRISPR-Cas9 genome editing, and suggest that therapeutic applications involving low-level Cas9 exposure will be particularly affected by chromatin status.

## Introduction

CRISPR-Cas9 is an RNA guided endonuclease involved in bacterial adaptive immunity, which has been repurposed as a highly efficient tool for eukaryotic genome editing [1–3]. In its natural form, Cas9 protein associates with a duplex of two RNA molecules: the crRNA, which recognises a short section of target DNA (the “protospacer”), and a tracrRNA, which acts as a scaffold to link the crRNA and Cas9 endonuclease. Most genome editing applications use a single guide RNA molecule (sgRNA) resulting from an engineered fusion of these two components. After target DNA cleavage, mutations arise through the action of cellular DNA repair pathways. Non-homologous end-joining (NHEJ, including both classical and microhomology-mediated pathways) can yield short insertions and deletions suitable for gene knockout, whereas homology-directed repair pathways utilise exogenous donor templates to introduce precise sequence changes.

It is well established that genetic properties of the genomic target site and sgRNA molecule have a significant effect on the efficiency of CRISPR mutagenesis [4–6]. However, Cas9, being prokaryotic in origin, did not evolve to cope with the complex chromatinised environment of the eukaryotic genome. Despite prior studies in this area [4,7–14], the extent to which epigenetic properties of the target site, including DNA and histone modifications, influence mutation frequency and DNA repair outcome remains incompletely understood. Stably positioned nucleosomes act as a barrier to Cas9 binding and function on synthetic chromatin fibres [7,8,11], and in vivo [7], yet catalytically dead (d)Cas9 can open previously inaccessible regions of chromatin [15,16]. It has been reported that some sgRNAs show reduced activity within heterochromatin whereas others do not [13,14]. The reasons behind this paradox are unclear, but presumably involve other experimental variables that modify the influence of chromatin on CRISPR activity. Furthermore, it is widely accepted that double strand break (DSB) repair is influenced by the chromatin environment in which DSBs arise [17–21], and DSB repair is central to the mechanism of genome editing [22,23]. However, it is unclear whether pre-existing epigenetic properties of the target site impact upon the specific sequence changes that arise following Cas9 cleavage.

Genomic imprinting is a natural epigenetic process in which either the maternal or paternally derived copy of a gene is transcriptionally silenced. Essential regulatory elements within imprinted domains called ‘imprinting control regions’ undergo differential CpG methylation in the male and female germline. This leads to the establishment of monoallelic domains of heterochromatin in the early embryo that are maintained throughout somatic development [24]. These imprinted alleles carry all known hallmarks of constitutive heterochromatin, including post-translational histone modifications (H3K9me3, H4K20me3, histone hypoacetylation) and heterochromatin binding proteins (HP1γ).

Genomic imprinting has provided numerous insights into mechanisms of transcriptional regulation [25–28]. Because active and silent alleles of imprinted loci have identical DNA sequence, chromosomal position and potential exposure to diffusible regulators, allele-specific chromatin modifications must be sufficient to account for their differential expression [29]. Based on this principle, we postulated that genomic imprinting could be used to provide new insights into the influence of chromatin modifications on targeted mutagenesis.

## Results

Mouse embryonic stem cell (mESC) lines were derived from male F1 hybrid blastocysts of inter-subspecies crosses between (C57BL6/J (B6) and the *Mus m. molossinus* inbred strain JF1 (Figure 1A). These cells are heterozygous for strain-specific single nucleotide polymorphisms (SNPs) [30], which serve as genetic markers that distinguish maternal and paternal chromosomes. To control for possible genetic effects on mutagenesis arising from SNPs, we derived mESCs from reciprocal crosses (B6 female × JF1 male (B×J), and JF1 female × B6 male (J×B)), and used both cell lines in parallel wherever possible.

We targeted three maternally imprinted CpG islands: *KvDMR1* (hereafter referred to as *KvDMR*, Figure 1B, Figure S1A), *Impact* (Figure S2A) and *Inpp5f_v2* (Figure S3A). Maternally imprinted loci were selected due to their greater epigenetic stability during ESC culture compared to paternally inherited marks [31]. To determine whether these loci had distinct epigenetic configurations on maternal and paternal alleles in B×J and J×B mESCs, we performed allele-specific DNAse-I hypersensitivity assays (Figure S1B, Figure S2B, Figure S3B), and allele-specific ChIP experiments for H3K9me3 and H4K20me3 (Figure 1C, Figure S1C, S1D, Figure S2C, S2D, Figure S3C, S3D). In each case, paternally derived alleles were substantially more sensitive to DNAse-I digestion, whereas maternal alleles were highly enriched for heterochromatin marks. Nonetheless, in specific instances such as *Inpp5f_v2* in B×J cells, loss of imprinting (LOI) was evident from incomplete allelic enrichment of histone modifications (Figure S3D), and incomplete depletion of paternal alleles by DNAse-I (Figure S3B). To quantify the degree of LOI, CpG methylation levels were assessed by PCR amplification of target sites from bisulfite-modified template DNA followed by high-throughput amplicon sequencing. As expected, loss of imprinting was observed to a variable degree in some, but not all cases, and at least one cell line maintained a substantial degree of imprinting at all three loci (Figure 1D, Figure S2E, Figure S3E). Instances where imprinted CpG methylation fell below 50% of expected levels were excluded from further analysis.

**Figure 1.**
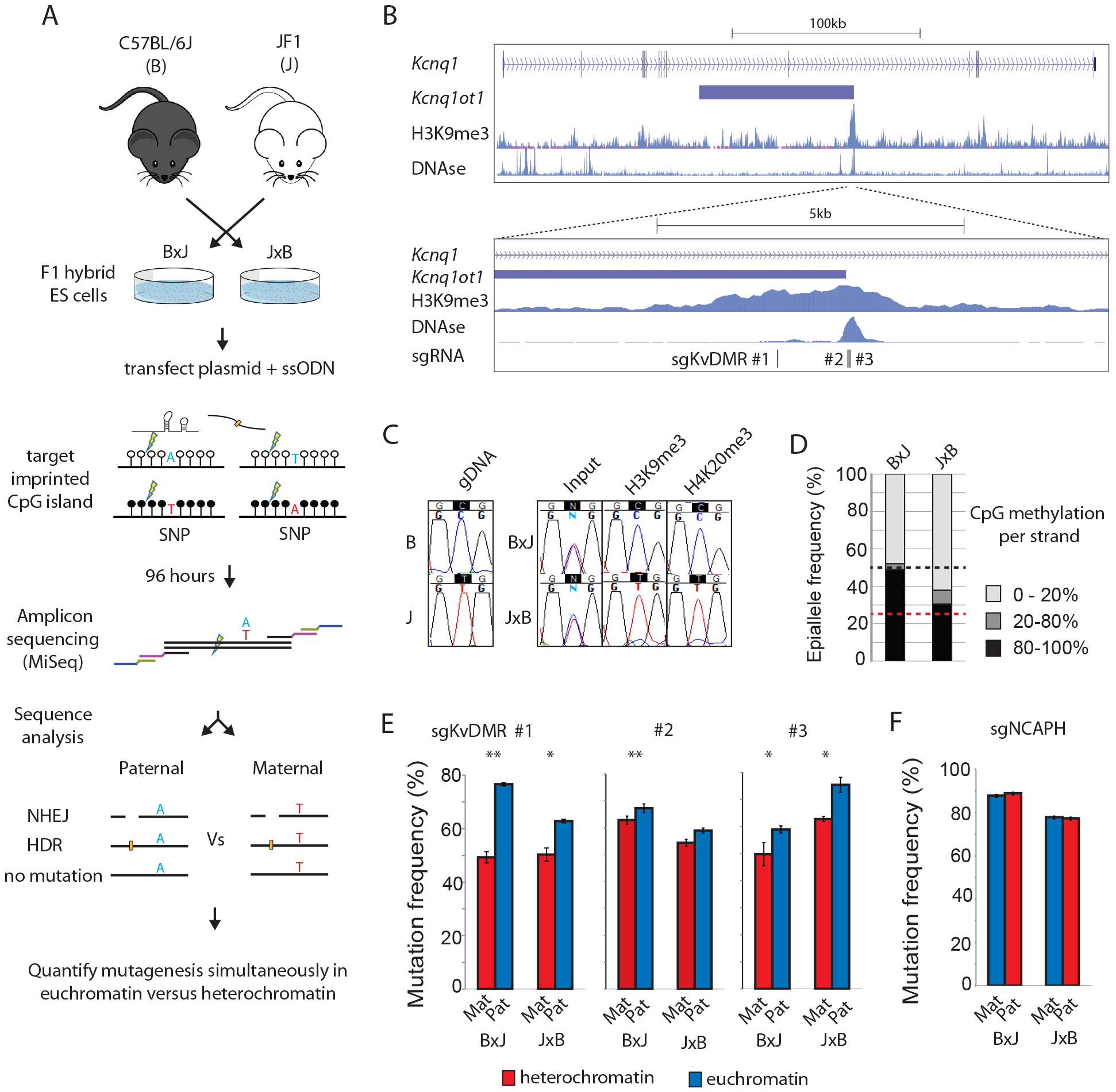
Imprinted chromatin as a model system to quantify epigenetic influences on genome editing. **A**. Schematic outlining the experimental workflow. Throughout the text, F1 hybrid cell lines are depicted with the maternal strain denoted before the paternal strain (ie. In B×J - B is maternal and J paternal). sgRNAs are designed to cleave approximately 40 −100bp from a heterozygous SNP (A/T) within imprinted chromatin (open and closed circles). MiSeq amplicons span both the SNP and site of mutation, which allows simultaneous assessment of genome editing outcome and parental allele at high-throughput. **B**. (top) Schematic showing the imprinted mouse *Kcnq1* gene including H3K9me3 ChIP and DNAse-I-seq data from mouse ESCs available through EncODE (ENCSR000CBH, GSM1014187) (bottom). Higher resolution view of the *KvDMR* imprinted CpG island within *Kcnq1*, showing the position of three sgRNAs used in panel E. **C.** Allele-specific enrichment of H3K9me3 and H4K20me3. PCR fragments spanning the target sites of sgKvDMR#2 & #3 were amplified from input, or ChIP DNA prior to Sanger sequencing across an allelic SNP. gDNA = genomic DNA from purebred mice. **D.** CpG methylation at the *KvDMR* locus. Bisulfite-converted genomic DNA was subjected to amplicon sequencing across a region spanning 13 CpG dinucleotides (Figure S1A), and reads were classified according to the proportion of non-converted (methylated) CpGs. The black dashed line indicates the expected level of methylation across all alleles when imprinting is completely maintained, and the red line the level with 50% loss of imprinting **E.** Allele-specific mutation frequencies for KvDMR sgRNAs #1 - 3. Error bars represent SEM of 3 biological replicates, p-values denote two-tailed paired t-tests of difference between maternal (Mat) and paternal (Pat) alleles. * p < 0.01, ** p < 0.001. **F.** Allele-specific mutation frequencies from experiments using an sgRNA (sgNCAPH) targeting a non imprinted locus, presented as in panel E.

We designed 3 different sgRNAs to target protospacer sequences within KvDMR (Figure 1B, Figure S1A). mESCs were transfected with Cas9 and sgRNA expressed from plasmid pX459v2 [32], together with a single stranded oligodeoxynucleotide (ssODN) donor template that introduced point mutations to prevent re-cutting following homology-directed repair (HDR). Transfected cells were selected in puromycin and collected as a pool 4 days after transfection. Editing was quantified by Illumina sequencing of PCR amplicons spanning both the site of cleavage and an allelic SNP (Figure 1A, Figure S1A, for detailed experimental protocols see Supplemental Methods). This allowed detailed assessment of mutagenic repair separately on maternal and paternal chromosomes, including the ratio of edits arising via either NHEJ or HDR pathways (see below).

We first compared the frequency of all edits (NHEJ+HDR) on maternal versus paternal alleles. All three sgRNAs yielded more mutations on the active paternal allele compared to the repressed maternal allele (Figure 1E), whereas a control, non-imprinted locus (*NCAPH*) showed no such allelic bias (Figure 1F). The effect of imprinted chromatin was remarkably subtle in this context: 1.2 – 1.6 fold, even in B×J cells where imprinting was completely maintained (Figure 1D).

To account for these results, we reasoned that CRISPR might less efficiently overcome the heterochromatin barrier when the intracellular concentration of Cas9 is low [33]. To test this hypothesis, KvDMR sgRNA#3 was expressed from plasmid pX458, in which spCas9 is fused to eGFP via a self-cleaving 2A peptide. eGFP levels therefore serve as a reporter of Cas9 translation (Figure 2A). Flow cytometry revealed that Cas9 translation levels were highly variable between cells at 24 hours post-transfection (Figure 2B). Cells were purified by fluorescent activated cell sorting (FACS) into three categories based on eGFP fluorescence and then collected either immediately (24h, Figure S4) or following a further 3 days in culture (Figure 2). Strikingly, BxJ cells expressing Cas9 at low levels showed a profound (5.3-fold) reduction in mutation frequency on the silent maternal compared to the active paternal allele after four days of exposure. At intermediate levels of Cas9-eGFP expression the mutational bias was moderate (2.6-fold), whereas high expression yielded only subtle differences between alleles (~1.2-fold)(Figure 2C). J×B cells showed the same trend but mutations on the maternal allele were more frequent, consistent with ~30% loss of imprinting (Figure 1D). Heterochromatin therefore impedes mutagenesis to a greater extent when the intracellular concentration of Cas9 is low.

**Figure 2:**
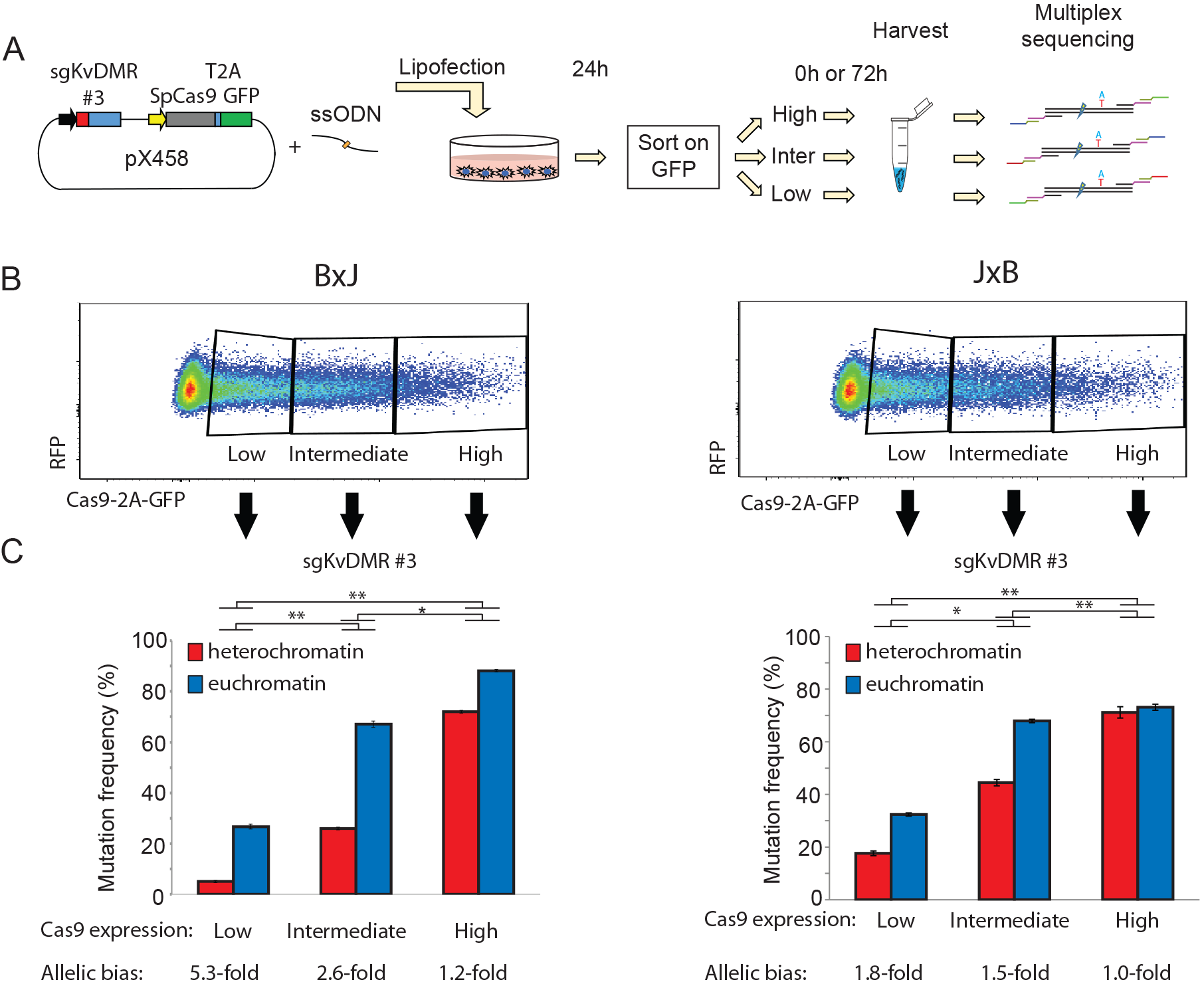
Heterochromatin impedes genome editing in a Cas9-concentration-dependent manner. **A.** Schematic outlining the experimental workflow. After FACS, cells were either harvested immediately (Figure S4) or cultured for a further 72 hours (this figure). **B**. Flow cytometry profiles show widely variable expression of Cas9-2A-eGFP at 24 hours following transfection with guide gKvDMR#3 (Figure 1B) expressed from pX458 (see panel A). **C.** Allele-specific mutation analysis within cell populations expressing different levels of Cas9, FACS-purified 24 hours post-transfection using the gating scheme in panel B and then cultured for a further 72 hours before harvesting. Allelic differences are less pronounced in JxB cells due to partial loss of imprinted heterochromatin on maternal alleles in this cell line (Figure 1D). Error bars represent SEM of 3 biological replicates. Asterisks denote p-values for unpaired t-tests on the fold-difference between maternal versus paternal allele mutation frequencies at different levels of Cas9-eGFP expression. * p < 0.01, ** p < 0.001.

Single particle tracking experiments have shown that the efficiency of target searching by catalytically dead (d)Cas9 is reduced within heterochromatin [9]. Whether this impacts upon mutagenesis with Cas9 nuclease was not tested. To determine whether heterochromatin delays mutation kinetics, we targeted the *Impact* imprinted locus (Figure S2A-E), using a highly active sgRNA (sgImpact) that yielded similar frequencies of mutation on maternal and paternal alleles after 4 days of exposure (Figure S2F). B×J cells were collected at 4 hour intervals following transfection and allele-specific mutagenesis was quantified as described above (Figure 3A). As expected, the frequency of mutations across both alleles increased steadily from 8 hours to 48 hours following transfection, but mutations were more skewed towards the active paternal allele at earlier compared to later time points (Figure 3B). Using sgRNAs targeting two additional imprinted loci (sgKvDMR#1 (Figure S1) and sgInpp5f_v2 (Figure S3)), we observed stronger skewing towards allelic target sites within euchromatin at early (16 hour) compared to later (96 hour) time points (Figure 3C, 3D). This effect was most striking in cells exposed to high concentrations of Cas9, for which a large majority (78%) of mutations present in euchromatin following 96 hours of exposure were found to occur within the first 24 hours (Figure 3E). Within heterochromatin, only 23% of mutations present at 96 hours had occurred by this earlier time point (Figure 3E). We conclude that heterochromatin impairs the kinetics of mutagenesis in a manner that depends on the level of intracellular Cas9 expression. However, target sites within heterochromatin ultimately reach similar frequencies of mutation upon sustained CRISPR exposure.

**Figure 3:**
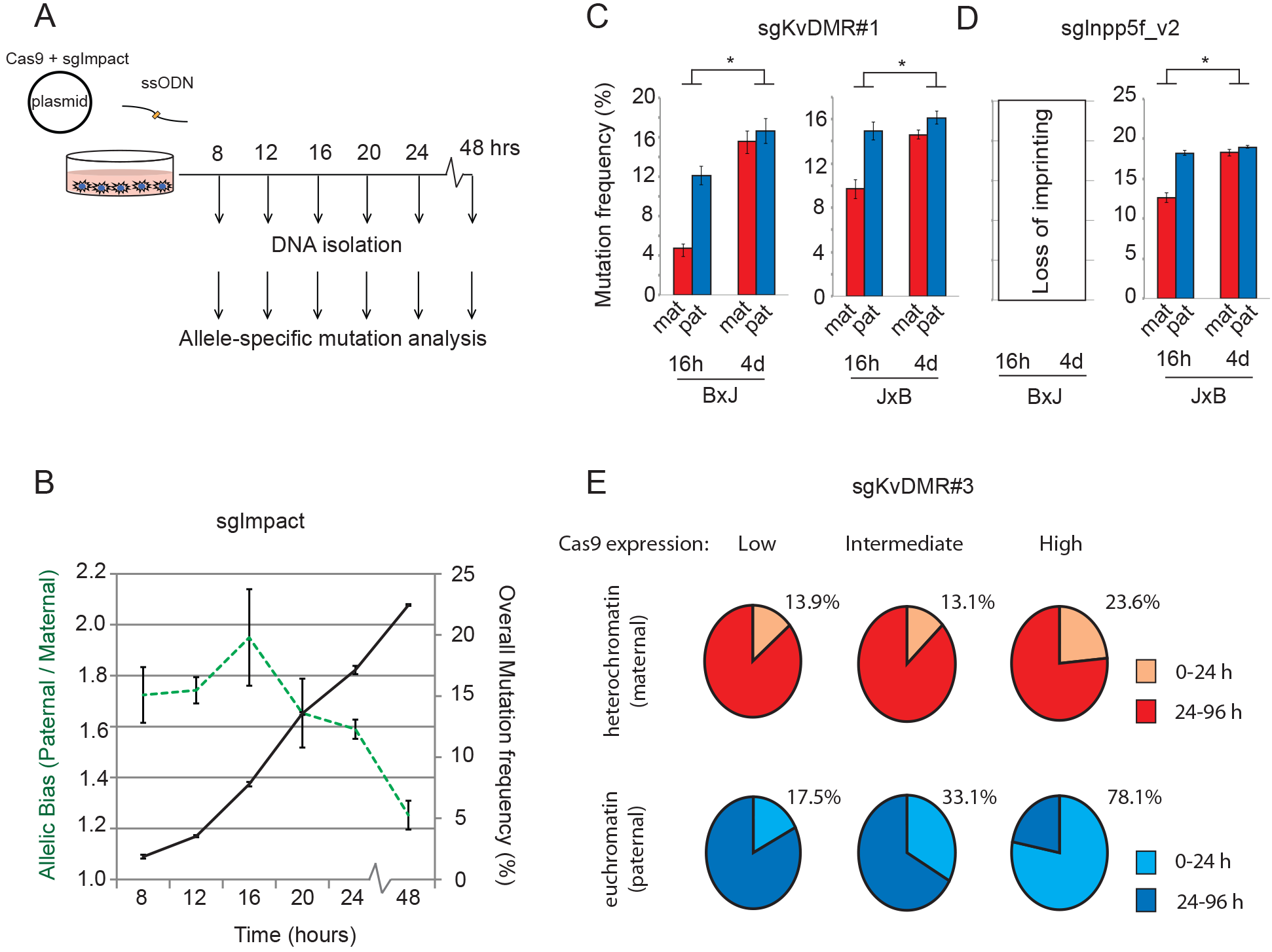
Heterochromatin impairs the kinetics of CRISPR mutagenesis. Schematic depicting the experimental workflow for the time-course experiment in panel B. **B.** Overlaid line graphs depict total mutation rates (black solid line, right y-axis) and skewing towards the euchromatic paternal allele (green dashed line, left y-axis) over time using sgImpact targeting the imprinted *Impact* locus (Figure S2) in B×J cells. Error bars represent SEM of 3 biological replicates. The short time frame prevented selection for transfected cells in this experiment. **C.** Allele-specific mutation frequencies at 16 hours (16h) and 4 days (4d) post-transfection for experiments using an sgRNA targeting KvDMR (sgKvDMR#1 – Figure S1A). Asterisks denote *p* < 0.05 for unpaired t-tests on the fold difference in maternal versus paternal allele mutation frequencies between timepoints. **D.** As above, using an sgRNA targeting the imprinted *Inpp5f_v2* promoter (Figure S3A). Note that a majority of maternal chromosomes had lost imprinting at this locus in B×J cells (Figure S3E), hence, only J×B data are shown. **E.** Pie charts show mutation frequencies observed 24 hours post transfection, expressed as a percentage of the mutation frequency in cells collected after 96 hours. Data are derived from the experiment described in Figures 2 and S4, with mutation frequencies broken down by both parental allele and Cas9 expression level. Experiments used sgKvDMR#3 in BxJ cells and were conducted in biological triplicate as described in Figure 2A, with cells collected either immediately after sorting on Cas9-2A-eGFP (24h) or after a further 72 hours in culture.

The repair of double strand breaks (DSBs) induced by Cas9-independent routes is thought to be influenced by the pre-existing chromatin environment at the site of cleavage [17,19–21]. However, whether DNA accessibility and/or epigenetic modification of DNA and histone proteins can influence the outcome of CRISPR mutagenesis, particularly the frequency of mutations arising via NHEJ versus HDR-mediated pathways, is not known. Imprinted genes provide an ideal system with which to address this question.

For 5 sgRNAs targeting imprinted heterochromatin, mutational profiles were calculated separately from sequencing reads originating from maternal (repressed) versus paternal (active) alleles (Figure 4A, Supplementary Methods). Surprisingly, no consistent allelic biases were evident in the NHEJ:HDR ratios at four days post-transfection (Figure 4B), but the rate of HDR varied by up to 3-fold between loci. This suggests that DNA sequence features of the target and HDR template molecules [5,34] are more important than epigenetic properties in determining HDR efficiency. Even at earlier time points, when overall mutation frequencies are significantly higher on paternal chromosomes (Figure 3B), the ratio of HDR to NHEJ was not significantly different on maternal and paternal alleles (Figure 4C). It is important to stress that our assay cannot, in its current form, measure non-mutagenic DSB repair that does not lead to genome edits. Nonetheless, the data suggest that the relative efficiency of sequence changes occurring via NHEJ or HDR is not substantially affected by the prior chromatin state. We note that a recent study in Drosophila found that DSB repair kinetics and pathway choice were similar in euchromatin versus heterochromatin following I-SceI cleavage [35].

**Figure 4:**
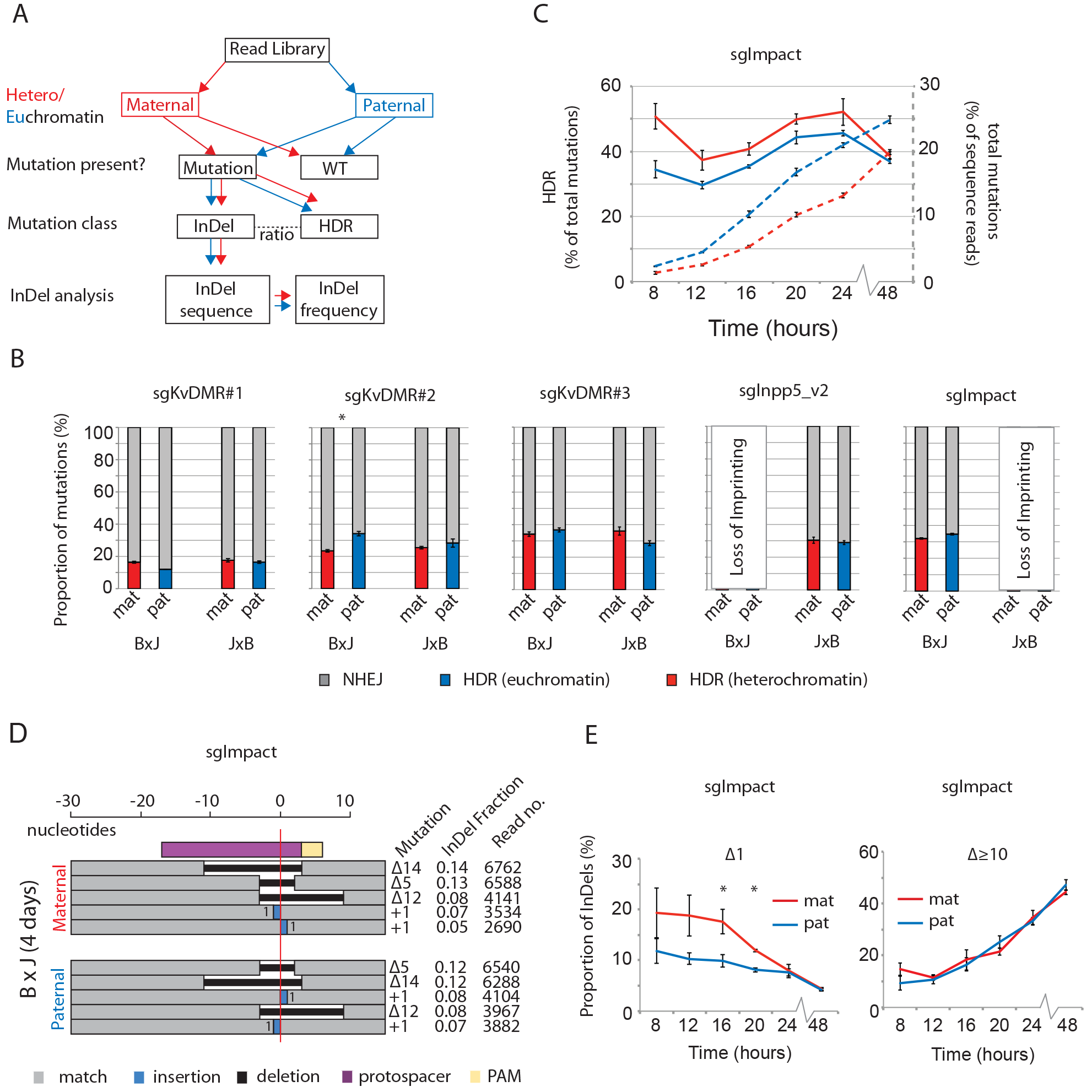
The efficiency of homology-directed repair is unaffected by heterochromatin. **A**. Schematic outlining the sequence analysis pipeline for allele-specific mutation analysis, described in full in the Supplemental Methods. **B.** The relative frequency of mutations arising from NHEJ versus HDR in cells collected at 4 days post-transfection, for 5 sgRNAs with target sites within imprinted heterochromatin (Figure S1A, S2A, S3A). Experiments at which imprinted CpG methylation fell below 50% of expected levels (Figure S2E, S3E) were excluded. Asterisk denotes bonferroni-corrected p-value of < 0.05 from paired t-tests of difference between HDR frequencies on maternal versus paternal alleles. **C.** Allele-specific proportions of mutations occurring via HDR (solid lines, left y-axis) over the time course experiment featured in Figure 3B. Allele-specific frequencies of all mutations (HDR + NHEJ) are shown as dashed lines (right y-axis) for comparison. Despite subtly higher frequencies of HDR on maternal alleles, p-values for paired t-tests were >0.05 at all timepoints. **D.** The size and frequency of the top 5 most common InDels 4 days following transfection with an sgRNA targeting the Impact locus (Figure S2A), broken down by parental allele. The horizontal red line denotes the predicted cleavage site. Deletion sizes are depicted against the scale bar at the top, and for insertions the number of inserted bases is indicated next to the blue rectangle. The fraction of Indels was calculated as the number of reads corresponding to each specific mutation expressed as a proportion of all InDel-containing reads (Supplementary Methods). **E.** Changes in the proportion of reads containing deletions of a single nucleotide (Δ1) or 10 or more nucleotides (Δ≥10) over a 48 hour time course experiment using Impact sgRNA in B×J cells (Figure 2A, 2B). Error bars represent SEM from 3 biological replicate experiments. Asterisks denote p-values < 0.05 for paired t-tests of difference between maternal and paternal alleles.

Next we asked whether chromatin modifications influenced the spectrum and frequency of different InDel mutation classes produced via non-homologous end-joining. In line with a recent large-scale deep sequencing study of InDels induced by Cas9 in cancer cell lines [23], we found that each sgRNA produced its own characteristic InDel pattern, with the top 5 recurrent mutations comprising 30 – 60% of all sequencing reads in cells collected 4 days following transfection (Figure 4D, Figure S5). Strikingly, the same mutations recurred on maternal and paternal chromosomes (Figure 4D, Figure S5) despite these allelic target sites starting in very different epigenetic states (Figure 1C, D, Figure S1, S2, S3).

In time course experiments, Van Overbeek *et al.* showed that the relative frequency of different InDel classes changed over time when Cas9 was continually expressed [23]. Our data confirmed that single nucleotide deletions tend to be more frequent at early time points, but larger deletions make up a progressively greater fraction of all InDels as the mutation profile matures (Figure 4E). This pattern likely stems from the susceptibility of single nucleotide deletions to repeated cycles of cleavage and repair until the sgRNA binding site is destroyed[23]. Although this trend was evident in both heterochromatin and euchromatin, single nucleotide deletions were more frequent on silenced compared to active alleles at earlier, but not late time points (Figure 4E). We note that overall mutation frequencies are lower in heterochromatin at these earlier time points (Figure 3). This might suggest that euchromatin provides an environment in which cycles of cleavage and repair can occur more rapidly, causing the InDel profile to mature at a quicker rate.

## Discussion

In this study, we have used the classical epigenetic model system of genomic imprinting to determine the effect of chromatin context on CRISPR-Cas9 genome editing. This internally controlled approach allowed us to identify key experimental parameters (intracellular Cas9 expression level and duration of exposure) that determine the extent to which repressed chromatin impairs mutagenesis. Our findings are consistent with and extend those of previous studies in this area. The inhibitory effect of nucleosomes on Cas9 binding and cleavage is well established [7,8,11], and the sgRNAs used in this study targeted regions of allele-specific DNAse hypersensitivity (Figure S1B, Figure S2B, Figure S3B). On hypersensitive alleles, nucleosome-DNA interactions are less stable due to chromatin remodelling activities associated with RNA Polymerase II transcription [36].

We propose that the dynamic nature of chromatin at these sites would provide more opportunities for Cas9 complexes to bind and cleave their targets per unit of time. Conversely, mutations accumulate more slowly in heterochromatin, where nucleosomes marked by H3K9me3 and H4K20me3 more effectively occlude Cas9 complexes. Mutagenesis still occurs within heterochromatin, albeit at a lower rate, presumably due to residual nucleosome breathing [11] and remodelling associated with DNA replication. Elevated concentrations of Cas9 increase the likelihood of mutation through mass action: an effect that we observed in both heterochromatin and euchromatin (Figure 2C), and which caused saturation to be reached more rapidly. Under optimal experimental conditions where no loss of imprinting occurred (Figure 1D), heterochromatin impeded mutagenesis by almost 7-fold (Figure S4).

In practical terms, our findings suggest that chromatin state is a particularly important consideration during procedures where the level of Cas9 exposure is kept low. This would be relevant in a clinical setting, where it is desirable to minimise exposure in order to avoid undesirable off-target mutations [37]. Indeed, our data support the use of epigenomic data for the prediction of off-target mutagenesis [38], and suggest that reducing Cas9 exposure would increase target specificity to the greatest extent when off-target sites are embedded within heterochromatin but on-target sites are within accessible regions.

We also addressed, to our knowledge for the first time, whether local chromatin state influences the relative frequency of CRISPR-Cas9 genome edits occurring via NHEJ versus HDR. We found that this important aspect of genome editing was not significantly different between heterochromatin and euchromatin. This is somewhat surprising in light of prior reports that chromatin modifications influence repair pathway choice in other contexts [19,20]. It is possible that chromatin remodelling events associated with Cas9 binding [15,16] are sufficient to overcome any prior differences in chromatin state, which might otherwise influence the outcome of DNA repair.

In summary, we show that allele-specific epigenetic model systems such as genomic imprinting can provide new insights into mechanisms of genome editing in a physiological setting. Given the expanding range of synthetic DNA binding proteins now used in research, biotechnology and medicine [39–43], this approach can provide further insights into their mode of interaction with chromatin *in vivo*. In the future, it will be of interest to extend this study to assess other chromatin states, such as transcribed versus non-transcribed imprinted gene bodies, and between targets on the active versus inactive X chromosome.

## Materials and Methods

### Cell culture and transfection

ESC lines were derived from male F1 hybrid blastocysts in 2i, serum-free medium using previously described methods[44]. ESCs were maintained on gelatin-coated dishes in ESGRO-complete-plus medium (Millipore, SF001-500P), under serum- and feeder-free conditions. All experiments, including mutagenesis and validation of allele-specific chromatin status, were performed on cells at passages 6 – 12. A modal chromosome number of 39 was confirmed by counting metaphase chromosomes of cells at passage 11. Protospacer sequences were selected using the online tool hosted by the Broad Institute (https://portals.broadinstitute.org/gpp/public/analysis-tools/sgrna-design), within three loci previously described in the literature to exhibit allele-specific CpG methylation[45]. In all experiments, both Cas9 and sgRNA were expressed from the same plasmid, which was transfected together with a 150 nucleotide single stranded oligonucleotide (ssOligo) which served as a template for homology-directed repair. ssOligos introduced nucleotide substitutions which removed the NGG PAM motif to prevent further cleavage. For the experiments presented in Figure 2 and Figure 3E, sgRNA and Cas9-2A-eGFP were expressed from plasmid backbone pX458, whereas all other experiments used plasmid backbone pX459v2[32]. Sequences of guides and donor oligonucleotides are listed in Table S2. Transfections were performed in duplex, i.e. each transfection mix contained two separate plasmids encoding sgRNA and ssODNs to target two loci simultaneously. Experiments examining the effect of Cas9 expression level on mutagenesis (Figure 2, Figure 3E) were the exception; here, plasmids were transfected individually.

Approximately 16 hours before transfection, 3 × 10^5^ cells were seeded in each well of a 6 well plate. Transfections were conducted using Lipofectamine 2000 (Invitrogen) according to the manufacturer’s protocol with the following modification: Transfection mix comprised a total of 3μg plasmid and 150ng oligonucleotide donor in 10 μl of P2000 reagent. Transfection efficiencies ranged from 15 – 50%. For all editing experiments that did not involve time points or Cas9-2A-eGFP selection, successfully transfected cells were selected in medium containing puromycin (1.6μg/ml) 24 hours following transfection. Puromycin was washed out together with dead cells at 48 hours following transfection, then genomic DNA was harvested from pooled cells at 96 hours. For the experiments in Figure 2 and 3E, cells were FACS purified using the gating strategy shown in Figure 2B at 24 hours following transfection. Each sorted population was split 50:50, with half harvested immediately and the remainder after a further 72 hours in culture. Transfected cells were not selected during any of the time course experiments presented in Figures 3 & 4. Sequences of guides and donor oligonucleotides are listed in Table S2.

### Locus specific amplification and MiSeq library preparation

DNA was isolated from edited cells using the DNeasy Blood and Tissue Kit (Qiagen) with RNAse treatment according to the manufacturer’s protocol. Each biological replicate used 50ng of template DNA, corresponding to 8,333 diploid genomes. Adaptors and barcodes necessary for multiplexed high-throughput amplicon sequencing were added using a two round PCR procedure. In the first round, locus-specific primers were designed to span regions encompassing both the editing site and an allelic SNP which allowed the origin of each sequence read to be traced to the maternal or paternal allele. First round primers contained 5’ extensions with a random hexamer, binding sites for illumina sequencing primers, and binding sites for universal primers necessary for the second round of cycling. Edited loci were amplified for 25 cycles using High Fidelity Phusion Polymerase (NEB). PCR products were purified using AMPure beads (Beckman Coulter) according to the manufacturer’s instructions and eluted in 50μl. 10μl of eluate was taken forward to a second round of PCR for 8 cycles. The second round of PCR used universal primers that contained unique indices based on the i5 and i7 sequences from the Nextera library prep kit (Illumina). This enabled multiplexing of libraries on a single flow cell. Locus specific and universal primer sequences are listed in Table S2. Amplified products were purified using AMPure beads, eluted in 25μl and then concentration and product size were verified on an Agilent Bioanalyser. Libraries were pooled at equimolar ratio and run on an Illumina MiSeq to obtain 150bp paired-end reads. Library details including read numbers are listed in Table S1.

### Bisulphite sequencing

DNA was purified from unedited control cells harvested at equivalent passage number to edited populations (8 – 12) using the DNeasy Blood and Tissue Kit (Qiagen). 0.5μg of DNA was subjected to bisulfite conversion using the EZ DNA methylation kit (Zymo) according to the manufacturer′s instructions. Each converted sample was eluted in a 10μl volume, of which 2μl was used as a PCR template. The generation of libraries for illumina sequencing proceeded as described above with one modification: the first round of PCR comprised 35 cycles rather than 25. A single library was generated for each locus.

### Chromatin Immunoprecipitation

The H3K9me3 ChIP-Seq track (GSM1000147) shown in Figure 1B and 1D is from the ENCODE mouse embryonic stem cell line BRUCE4 (C57BL/6J strain), visualised using the UCSC genome browser on GRC37/mm9. All ChIP assays presented in Figures 1, S1, S2 and S3 were performed on the hybrid lines used for mutagenesis studies. H3K9me3 (07-442, batch 2664282) and H4K20me3 (07-643, batch 2586586) antibodies used in ChIP experiments were purchased from Millipore. Approximately 10 million cells were harvested at approximately 80% confluency, trypsinised and washed in ice cold Phosphate Buffered Saline (PBS). Following centrifugation at 500.g, cells were resuspended in 1ml of ice cold NBA buffer (85mM NaCl, 5.5% sucrose, 10mM Tris-HCl pH 7.5, 0.2mM EDTA, 0.2mM PMSF, 1mM DTT, protease inhibitors). 1ml of NBB buffer (NBA buffer with 0.1% NP-40) was added, cells were incubated for 5 minutes on ice then centrifuged at 1000.g for 5 minutes at 4°C. The pellet was resuspended in 200μl of NBR buffer (85mM NaCl, 5.5% sucrose, 10mM Tris-HCl pH 7.5, 3mM MgCl2, 1.5mM CaCl2, 0.2mM PMSF, 1mM DTT) and centrifuged for a further 5 minutes at 4°C, then resuspended in 600ul NBR buffer. 10ul of RNAseA (10mg/ml) was added and incubated for 5 minutes at room temperature. 40 Boehringer units of MNAse (Sigma) were added, mixed by pipetting and incubated at 20°C for 10 minutes, with a further mix by pipetting after 5 minutes. Digestion was stopped by adding 600ul of MNase stop buffer (215mM NaCl, 10mM TrisHCl pH8, 20mM EDTA, 5.5% sucrose, 2% TritonX100, 0.2mM PMSF, 1mM DTT, 2× PMSF) and samples were stored at 4°C overnight.

40ul of protein A dynabeads (Invitrogen) were used per sample. After prewash in block solution (0.5% BSA in PBS), beads were mixed with 2.5ug antibody in 1ml block solution, incubated for 2 hours on a rotating wheel at 4°C and then washed in 200ul block solution. Chromatin was centrifuged at 13000RPM for 5 minutes at 4°C, and the supernatant transferred to a fresh tube with 10% set aside for use as input. 1ml of supernatant was added to the antibody bound beads together with 5ul of BSA (5mg/ml), before incubation at 4°C for 3 hours on a rotating wheel.

Three washes with ChIP-W1 buffer (150mM NaCl, 10mM Tris HCL pH8, 2mM EDTA, 1% NP40, 1% Sodium Deoxycholate) were performed in 1ml volume on a rotating wheel for 10 minutes at 4°C, followed by 1 wash in TE Buffer at room temperature without rotation. After the last wash beads were resuspended in 100ul of elution solution (0.1mM NaHCO_3_, 1% SDS), vortexed briefly and incubated at 37°C in a shaking thermomixer at 700rpm. The pH was adjusted to pH8 by adding 7ul of 2M Tris-Hcl pH6.8. Dynabeads were removed and the remaining solution (and input samples) was treated with 20ug of proteinase K for 1 hour at 55°C. ChIP and input DNA were purified on Qiagen PCR purification columns.

For relative quantification of ChIP DNA by real-time qPCR, DNA isolated from 10% of total MNase digested native chromatin was used to generate a standard curve (fivefold dilutions, from 10-0.08% total input) for IP samples. qPCR was performed in triplicate using SYBR Select mastermix (Applied Biosystems) on a LightCycler 480 II (Roche) with thermal cyling as follows: Initial Cycle 50°C for 2 minutes, 95°C for 2 minutes, then 40 cycles of 95°C for 15 seconds, 60°C for 50 seconds, 60°C for 10 seconds with a single acquisition. 0.5uL input or ChIP DNA was used in a total reaction volume of 20uL. For allele-specific enrichment analysis, regions spanning an allelic SNP were amplified using GoTaq (Promega), and amplicons were purified using AMPure beads and then subjected to Sanger sequencing. Primer sequences are listed in Table S2.

### DNase-I accessibility assay

DNAseI digestion was performed using a published protocol[46] with the following modifications. 20 × 10^6^ cells were trypsinised and resuspended in 5mL buffer A (15 mM Tris HCl (pH 7.6), 60 mM KCl, 15 mM NaCl, 1 mM EDTA, 0.5 mM EGTA, 0.5 mM spermidine, 0.15 mM spermine). Cells were lysed in the presence of 0.5 % (v/v) NP40, and nuclei were collected by centrifugation (2000g/5 minutes) and resuspended in 1mL digestion buffer (buffer A supplemented with 3 mM CaCl2, 75 mM NaCl). Digestions were carried out at 37 °C with 0-60 units of DNaseI (Sigma) per 100μL nuclei, for five minutes before the reaction was stopped by the addition of an equal volume of stop buffer (0.1 M NaCl, 0.1 % (w/v) SDS, 50 mM Tris-HCl (pH 8.0), 100 mM EDTA). The samples were treated with 2μg proteinase K at 55C overnight and DNA was recovered after extraction with phenol/chloroform and precipitation in ethanol. The DNA was then resuspended in TE buffer (10 mM Tris-HCl (pH 8.0), 1 mM EDTA), and concentration was measured using fluorometric quantitation (Qubit). Digested DNA was amplified for 30 cycles across regions containing an allelic SNP. Amplicons were purified using AMPure beads and then subjected to Sanger sequencing across regions of 300-600bp spanning an allelic polymorphism. Primer sequences

### Analysis of high throughput sequencing data

*All samples*. Reads were de-multiplexed and duplicate read pairs removed by FastUniq v1.1[47], and adaptors trimmed with TrimGalore v0.4.1 (https://www.bioinformatics.babraham.ac.uk/projects/trimgalore/). *Genomic sequencing of edited samples*. Trimmed and de-duplicated read pairs were aligned to mouse genome build GRCm38 using BWA v0.7.12[48]. Read pairs were extracted by the expected genomic region for each experiment, and assigned to the C57BL/6J or JF1 chromosome based on nucleotide identity at known polymorphic SNPs (http://molossinus.lab.nig.ac.jp/msmdb/index.jsp). Read pairs containing mutations originating from HDR were identified based on the expected sequence changes introduced from the oligonucleotide donors (Table S2); whereas read pairs containing insertions and deletions within 10bp of the cleavage site were identified as originating from fragments which had undergone NHEJ. Read pairs with evidence of neither were labelled as wild type. Indel length and type (insertion or deletion) were extracted from the NHEJ read pairs via a custom Perl script.

*Bisulfite sequencing of unedited samples*. Trimmed and de-duplicated read pairs were aligned to the bisulfite conversion indexed mouse genome build GRCm38 using Bismark v0.16.3[49] with Bowtie v2.2.6[50]. Read pairs that did not align were then separated and each end of the pair aligned as single end reads. The three resulting alignments were merged. Read pairs were extracted by the expected genomic region for each experiment,and assigned to the C57BL/6J or JF1 chromosome based on nucleotide identity at known polymorphic SNPs. The number of methylated CpGs in each read pair was counted using a custom Perl script examining the XM tag for each read in the relevant BAM file. All sequencing data have been deposited in the Sequence Read Archive under Study Accession SRP126405.

## Acknowledgements

We thank Feng Zhang’s laboratory for sharing the CRISPR plasmids used in this study through Addgene, Edinburgh Genomics for high throughput sequencing, and the IGMM Flow Cytometry Facility for FACS. We are also grateful to Wendy Bickmore and Rebecca Holmes for comments on the manuscript, and to Nick Gilbert for useful discussions. AW’s laboratory is funded by a Sir Henry Dale Fellowship from the Wellcome Trust and Royal Society (102560/Z/13/Z). RF Acknowledges grant funding from the Fondation Recherche Medicale (FRM, grant DEQ20150331703).

---

**Figure S1:**
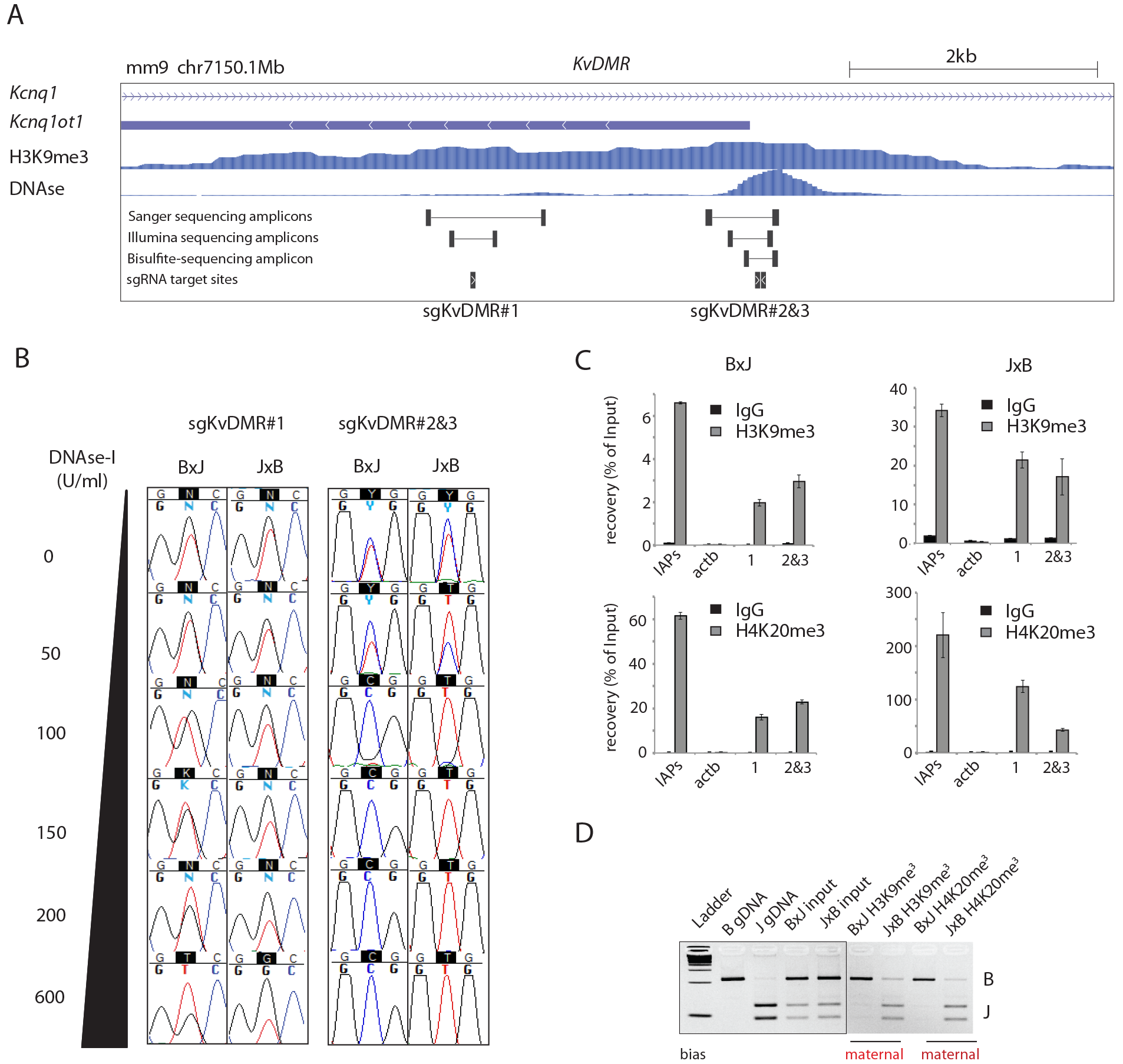
Allele-specific chromatin states at the imprinted *KvDMR* locus. **A.** UCSC screen drop showing the *KvDMR* locus, including the transcriptional start site for the *Kcnq1ot1* non-coding RNA, which is active from the paternal allele. H3K9me3 ChIP and DNAse-I-seq data from mESCs are available through EncODE (ENCSR000CBH, GSM1014187). Positions of sgRNA target sites and PCR amplicons used during the analysis are indicated. **B.** Allele-specific DNAse-I sensitivity of regions indicated in panel A. Note that Target 2 is within an annotated DNAse-I hypersensitive site whereas Target 1 is not. mESC nuclei were subjected to digestion with increasing concentrations of DNAse-I for 5 minutes at 37°C, before DNA extraction and Sanger sequencing across SNPs to reveal allele-specific differences in digestion at the regions indicated in panel A. **C.** Native ChIP enrichment for H3K9me3 and H4K20me3 marks at regions corresponding to sgRNA target site 1, and 2&3 (amplicons indicated in panel A). Enrichments are expressed relative to input, and error bars represent SEM of 3 technical replicates. qPCR primers spanning Intracisternal A particle (IAP) retrotransposons and the actb promoter serve as positive and negative controls, respectively. **D.** Allele-specific enrichment in ChIP DNA for the Target 1 region shown in panel A determined by RFLP analysis. The data are representative of two biological replicates for each mESC line.

**Figure S2:**
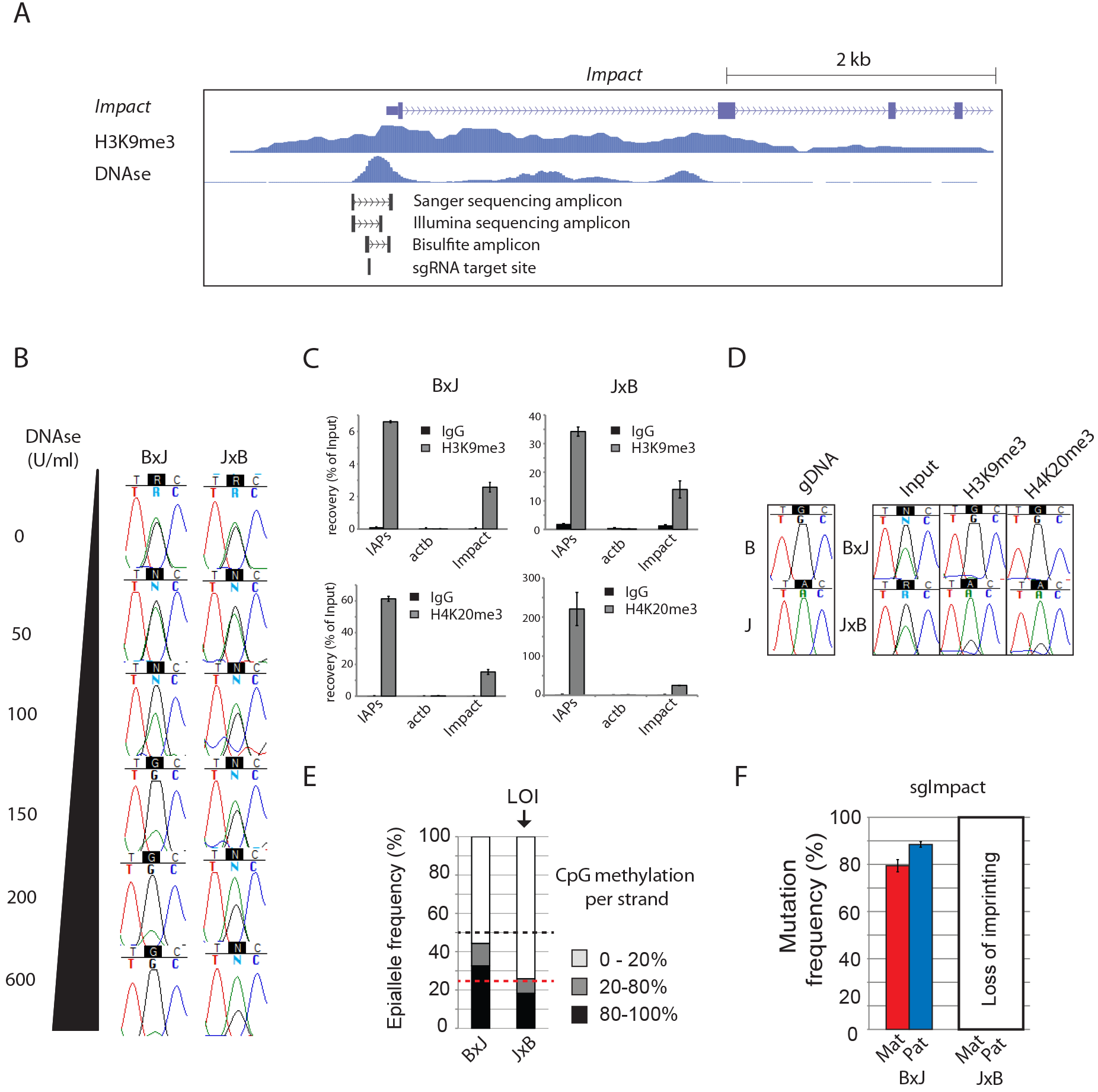
Allele-specific chromatin states at the imprinted *Impact* locus. **A**. UCSC screen drop showing the transcriptional start site for the *Impact* gene, which is active from the paternal allele. H3K9me3 ChIP and DNAse-I-seq data from mESCs are available through EncODE (ENCSR000CBH, GSM1014187). Positions of the sgRNA target site and PCR amplicons used during the analysis are indicated. **B.** Allele-specific DNAse-I sensitivity for a region spanning the target site, as indicated in panel A. **C.** ChIP enrichment for H3K9me3 and H4K20me3 marks at the *Impact* sgRNA target site. Enrichments are presented in the same manner as Figure S1C. **D.** Allele-specific enrichment of ChIP DNA at the *Impact* sgRNA target site determined by Sanger sequencing from ChIP DNA across an allelic SNP. ChIP experiments are representative of two biological replicates for each mESC line. **E.** CpG methylation at the *Impact* promoter presented as described for Figure 1D. The black dashed line indicates the expected level of methylation across all alleles when imprinting is completely maintained, and the red line the level with 50% loss of imprinting (LOI). **F.** Allele-specific mutation analysis from experiments using sgImpact in cells collected 4 days post-transfection. Data are presented as described in Figure 1E. Error bars depict SEM, n=3 biological replicates.

**Figure S3:**
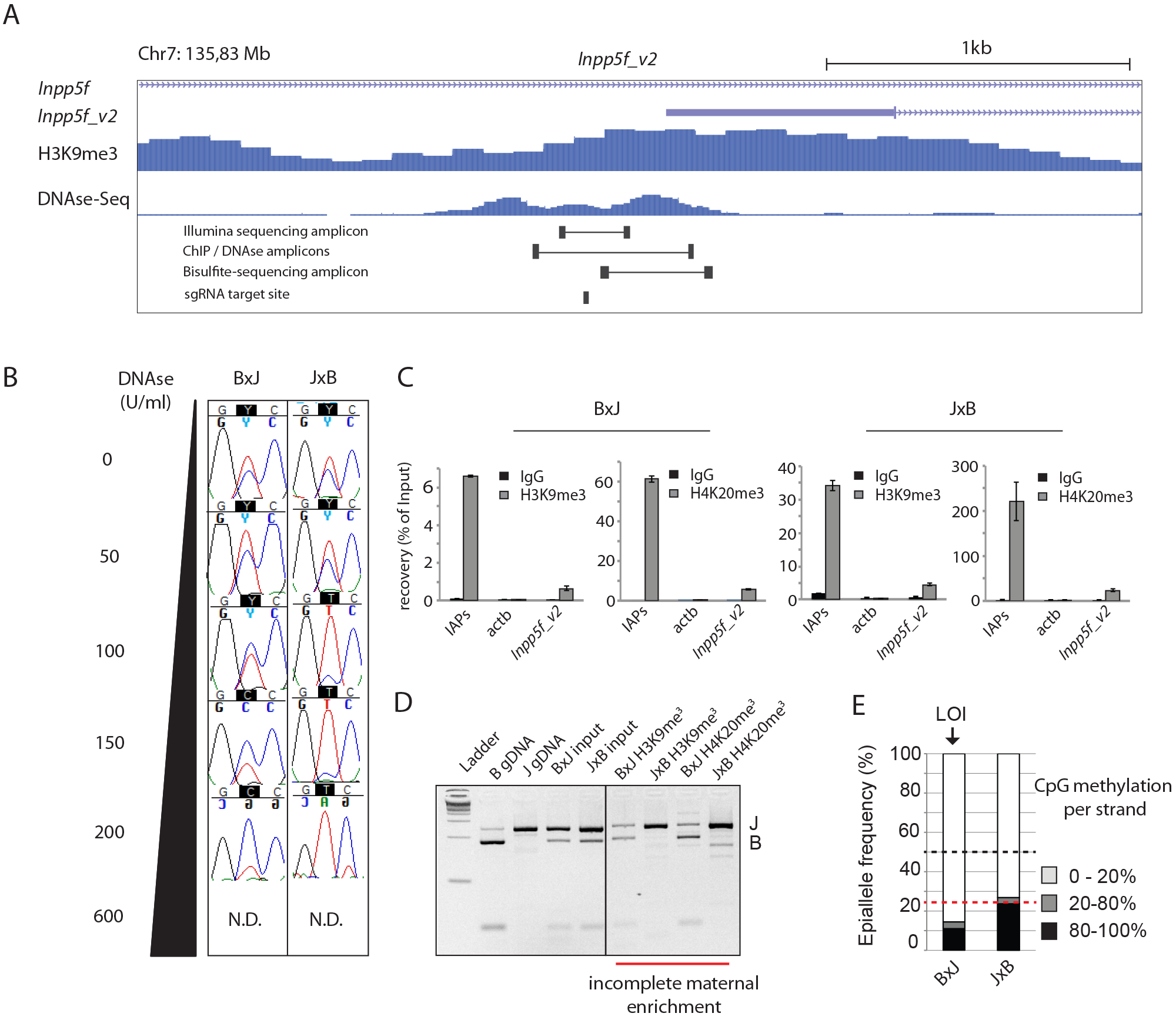
Allele-specific chromatin states at the imprinted *Inpp5f_v2* locus. **A**. UCSC screen drop showing the transcriptional start site for the *Inpp5f_v2* transcript, which initiates from the paternal allele. H3K9me3 ChIP and DNAse-I-seq data from mouse ESCs available through EncODE (ENCSR000CBH, GSM1014187). Positions of the sgRNA target site and PCR amplicons used during the analysis are indicated. **B.** Allele-specific DNAse-I sensitivity for a PCR amplicon spanning the *Inpp5f_v2* sgRNA target site, as described in Figure S1B. ND = not done due to poor PCR amplification in these samples. **C.** ChIP enrichment for H3K9me3 and H4K20me3 marks at the *Impact* sgRNA target site. Enrichments are presented in the same manner as Figure S1C. **D.** Allele-specific enrichment in ChIP experiments at the *Inpp5f_v2* sgRNA target site determined by RFLP analysis of PCR products amplified from ChIP DNA. ChIP experiments are representative of two biological replicates for each mESC line. **E.** CpG methylation at the *Impact* promoter presented as described for Figure 1D. The black dashed line indicates the expected level of methylation across all alleles when imprinting is completely maintained, and the red line the level with 50% loss of imprinting. Note the partial loss of imprinting which is evident in panels B, D and E, particularly in the B×J mESC line.

**Figure S4:**
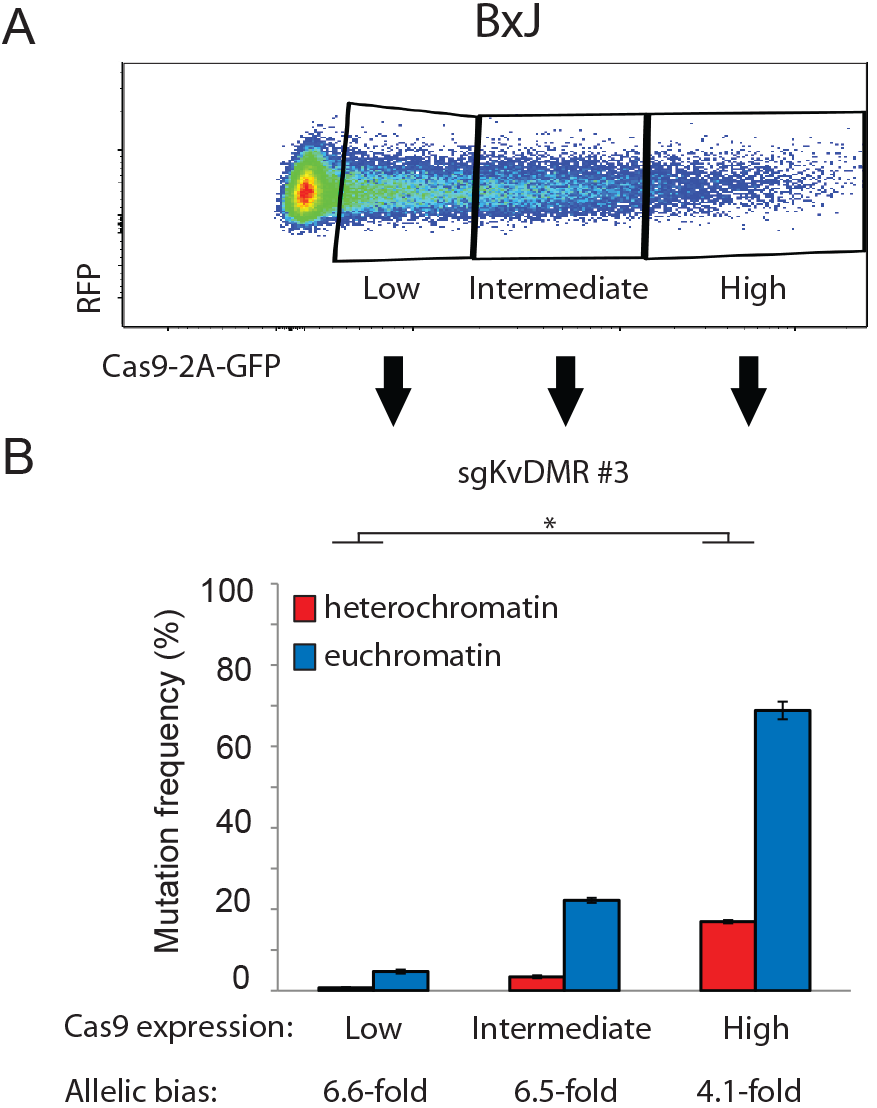
Heterochromatin impedes mutagenesis in a Cas9-concentration-dependent manner. **A.** B×J cells from the transfection shown in Figure 2A were FACS purified according to the gating scheme shown. Note that this panel depicts the same data shown in panel 2A. **B.** Allele-specific mutation analysis within cell populations expressing different levels of Cas9, as shown in panel A, FACS-purified 24 hours post-transfection and then subjected to allele-specific mutation analysis immediately, without further time in culture. Insufficient J×B cells were obtained following FACS to assess mutagenesis after 24h. Error bars represent SEM of 3 biological replicates. Asterisks denote p-values for unpaired t-tests on the fold-difference between maternal versus paternal allele mutation frequencies at different levels of Cas9-eGFP expression. * p < 0.05.

**Figure S5:**
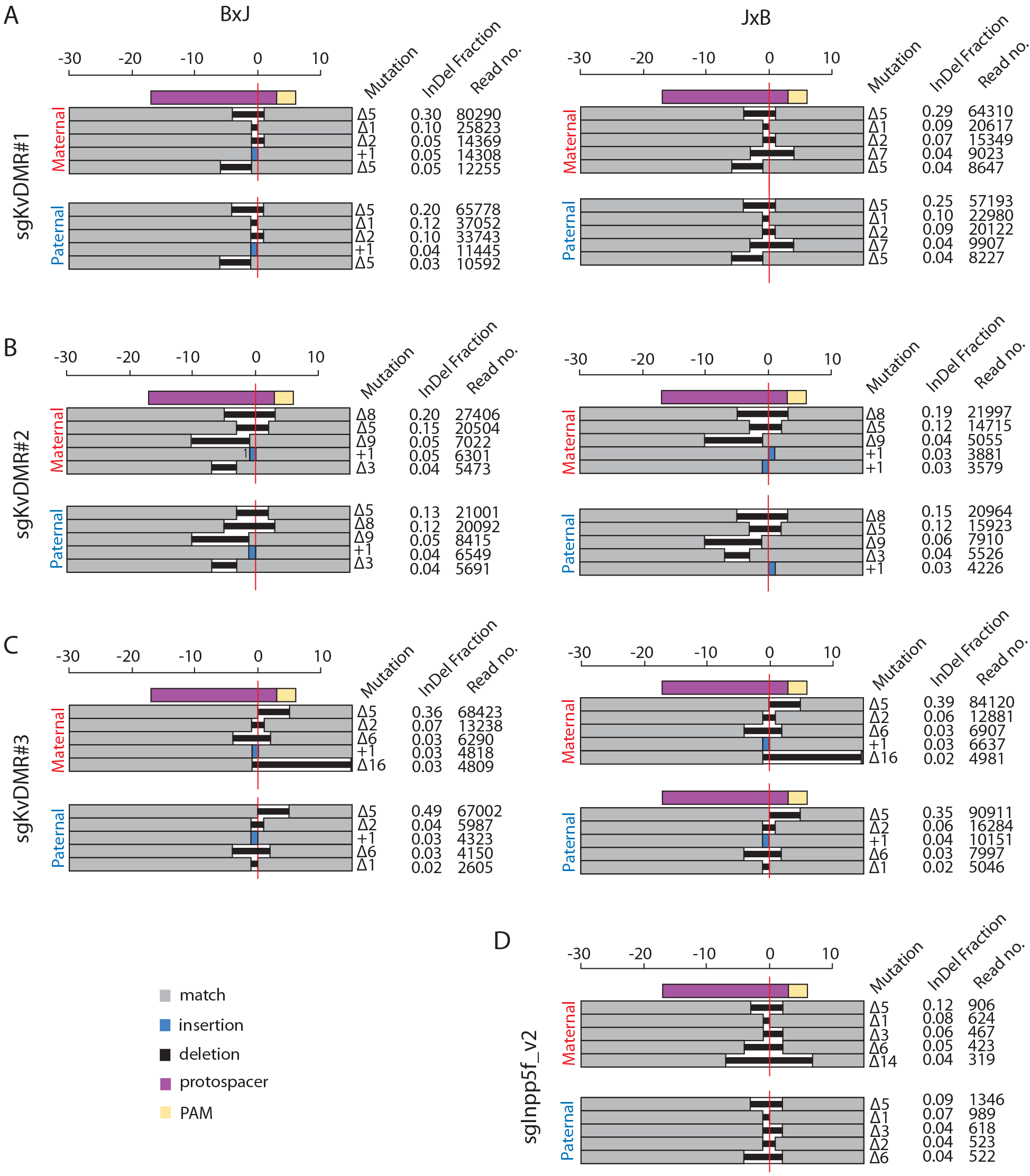
The same InDel classes recur in heterochromatin and euchromatin 4 days post-transfection. The size and frequency of the top 5 most common InDels (broken down by parental allele) produced by 4 different sgRNAs targeting imprinted heterochromatin. Edited genomic DNA was extracted 4 days following transfection with sgRNAs targeting the *KvDMR* (A, B, C) or *Inpp5f_v2* (D) imprinted loci in B×J (left) and J×B (right) cells. Note that a majority of maternal chromosomes had lost imprinting at the *Inpp5f_v2* locus in B×J cells (Figure S3E), hence, only J×B data are shown. Deletion sizes are depicted against the scale bar at the top of each panel, and the number of inserted bases is indicated next to the blue rectangle. The horizontal red line denotes the predicted cleavage site, and the colour key for all panels is situated at the bottom left of the figure. The fraction of Indels was calculated as the number of reads corresponding to each specific mutation class, expressed as a proportion of all InDel-containing reads (Supplementary Methods).

